# Is there a problem with methods skills in cognitive neuroscience? Evidence from an online survey

**DOI:** 10.1101/329458

**Authors:** Olaf Hauk

## Abstract

Cognitive neuroscience increasingly relies on complex data analysis methods. Researchers in this field come from highly diverse scientific backgrounds, such as psychology, engineering and medicine. This poses challenges with respect to acquisition of appropriate scientific computing and data analysis skills, as well as communication among researchers with different knowledge and skills sets. Are researchers in cognitive neuroscience adequately equipped to address these challenges? Here, we present evidence from an online survey of methods skills. Respondents (n=305) mainly comprised students and post-doctoral researchers working in the cognitive neurosciences. Multiple choice questions addressed a variety of basic and fundamental aspects of neuroimaging data analysis, such as signal analysis, linear algebra, and statistics. We analysed performance with respect to the following factors: undergraduate degree (grouped into Psychology, Methods, Biology), current researcher status (undergraduate student, PhD student, post-doctoral researcher), gender, and self-rated expertise levels. Overall accuracy was 72%. Not surprisingly, the Methods group performed best (87%), followed by Biology (73%) and Psychology (66%). Accuracy increased from undergraduate (59%) to PhD (74%) level, but not from PhD to post-doctoral (74%) level. The difference in performance for the Methods versus non-methods (Psychology/Biology) groups was particularly striking for questions related to signal analysis and linear algebra, two areas especially relevant to neuroimaging research. Self-rated methods expertise was not strongly predictive of performance. The majority of respondents (93%) indicated they would like to receive at least some additional training on the topics covered in this survey. In conclusion, methods skills among junior researchers in cognitive neuroscience can be improved, researchers are aware of this, and there is strong demand for more skills-oriented training opportunities. We hope that this survey will provide an empirical basis for the development of bespoke skills-oriented training programmes in cognitive neuroscience institutions.

## Introduction

Cognitive neuroscientists use physical measurements of neural activity and behaviour to study information processing in mind and brain. The development of novel experimental methodology has been a strong driving force behind psychological research in general (Greenwald, 2012). In recent decades, neuroimaging techniques such as functional magnetic resonance imaging (fMRI) and electro-/magnetoencephalography (EEG/MEG) have become standard tools for cognitive (neuro-)scientists. These techniques produce large amounts of data reflecting changes in metabolism or electrical activity in the brain (Huettel, Song, & McCarthy, 2009; Supek & Aine, 2014). Neuroimaging data can be analysed in a myriad of ways, involving basic pre-processing (e.g. filtering and artefact correction), model fitting (e.g. general linear models, structural equation modelling), and statistical analysis (e.g. t-tests, permutation tests, Bayes factor) (see e.g. https://en.wikibooks.org/wiki/SPM). Researchers involved in this work come from diverse backgrounds in psychology, cognitive science, medicine, biology, physiology, engineering, computer science, physics, mathematics, etc. This poses challenges with respect to the acquisition of the appropriate scientific computing and data analysis skills, as well as for communication among researchers from very different backgrounds.

A number of neuroimaging software packages offer standardised analysis pipelines for certain types of analysis problems (e.g. http://nipype.readthedocs.io, http://automaticanalysis.org), such as standard event-related fMRI or resting-state data, or conventional event-related potentials. Does this mean that empirical scientists no longer have to understand how their analysis pipelines work? There are several reasons why this is not the case.

First, few experiments are ever purely “standard” or “conventional”. A new research question, or a small change in a conventional experimental paradigm, may require a significant change of the analysis pipeline or parameter choices. The application of a novel, non-established, high-level analysis method is often the highlight of a study, e.g. in the case of multivariate pattern information analysis or graph-theoretical methods. It may not be possible to apply these methods in the same way as in similar previous studies, because there aren’t any.

Second, even for relatively standard analysis problems, there is often more than one solution, depending on how many people you ask. It is not uncommon to ask the nearest methods expert for advice, but it can be frustrating for a beginner to hear exactly the opposite advice from the next expert, especially if it is the reviewer of a paper or grant. One contributing factor to recently reported “replication crises” in cognitive science (Ioannidis, 2005; Open-Science-Collaboration, 2015) may be the large uncertainties in analysis procedures. They may lead to unintended or deliberate explorations of the parameter space (data mining or fishing). Even apparently simple issues relating to data analysis, such as “double dipping” or how to infer statistical interactions, are discussed in high-ranking publications (Kriegeskorte, Simmons, Bellgowan, & Baker, 2009; Nichols & Poline, 2009; Nieuwenhuis, Forstmann, & Wagenmakers, 2011; Vul, Harris, Winkielman, & Pashler, 2009). It should be good research practice to discuss any uncertainties in parameter choices and the implications these may have for interpretation of their findings or for future research.

Third, even researchers who do not employ high-level analysis methods or who are not involved in neuroimaging projects themselves need a certain level of understanding of data analysis methods in order to interpret the literature. The inferential chain from measurement through analysis to models of information processing in mind and brain can be complex (Carandini, 2012; Coltheart, 2013; Henson, 2005; Mather, Cacioppo, & Kanwisher, 2013; Page, 2006; Poldrack, 2006). Even a ubiquitous term such as “brain activation” can mean very different things in different contexts (Singh, 2012). The same holds for popular concepts such as “connectivity”, “oscillations”, “pattern information”, and so forth (Bastos & Schoffelen, 2015; Diedrichsen & Kriegeskorte, 2017; Friston, 2011; Haxby, 2012).

It will not be difficult to convince most cognitive neuroscientists that good methods skills are advantageous and will make their research more efficient. The greater their skills set, the greater their options and opportunities. At the same time, methods skills are clearly not the only skills required in cognitive neuroscience, raising the question of how much time a researcher should spend honing them relative to time spent on cognitive science, computational modelling, physiology, anatomy, etc. There cannot be a one-fits-all answer, as it depends on the goals and research environment of the individual researcher. However, in order to address this question, one can try to break the bigger problem down into several smaller ones.

In a first step, we can find out whether there is a problem at all, i.e. whether cognitive neuroscientists already have the basic skills required to understand the most common neuroimaging analysis methods. If there is room for improvement, then in a second step we can determine specifically where improvement is most needed, e.g. for which type of researcher and which type of methods skills. And finally, once we know where improvement is needed, we can find out whether researchers are aware of this and what opportunities there are to do something about it.

Surprisingly, there is currently no empirical basis for answering these questions. Because researchers enter the field from many different backgrounds, there is no common basic training for all researchers. Students from an engineering or physics background will have advanced skills in physical measurement methods, mathematics, signal processing and scientific computing, but may lack a background in statistics (as well as cognitive science and behavioural experimental techniques, which are not within the scope of this survey). Students of medicine and biology may have had basic training in physics and biostatistics, but not in scientific computing and signal processing. Students in psychology and social sciences usually have training in statistical methods, but not in signal processing and physical measurement methods. It is therefore likely that many of these researchers, whatever their backgrounds, start their careers in cognitive neuroscience with an incomplete methods skills set. Neuroimaging training programmes vary considerably across institutions. It is therefore unclear to what degree junior researchers acquire appropriate methods skills for cognitive neuroscience at different stages of their academic development.

Here, we evaluated the level of methods skills for researchers at different stages of their careers, mostly at post-graduate and post-doctoral level. We present results from an online survey of methods skills from 305 participants, mostly students and post-doctoral researchers working in the cognitive neurosciences. Questions in the survey were chosen to cover basic aspects of data acquisition and analysis in the cognitive neurosciences, with a focus on neuroimaging research (Figure 1). We report results broken down by undergraduate degree (grouped into Psychology, Methods, Biology), current researcher status (undergraduate, PhD student, post-doctoral researcher), and gender.

**Figure 1:**
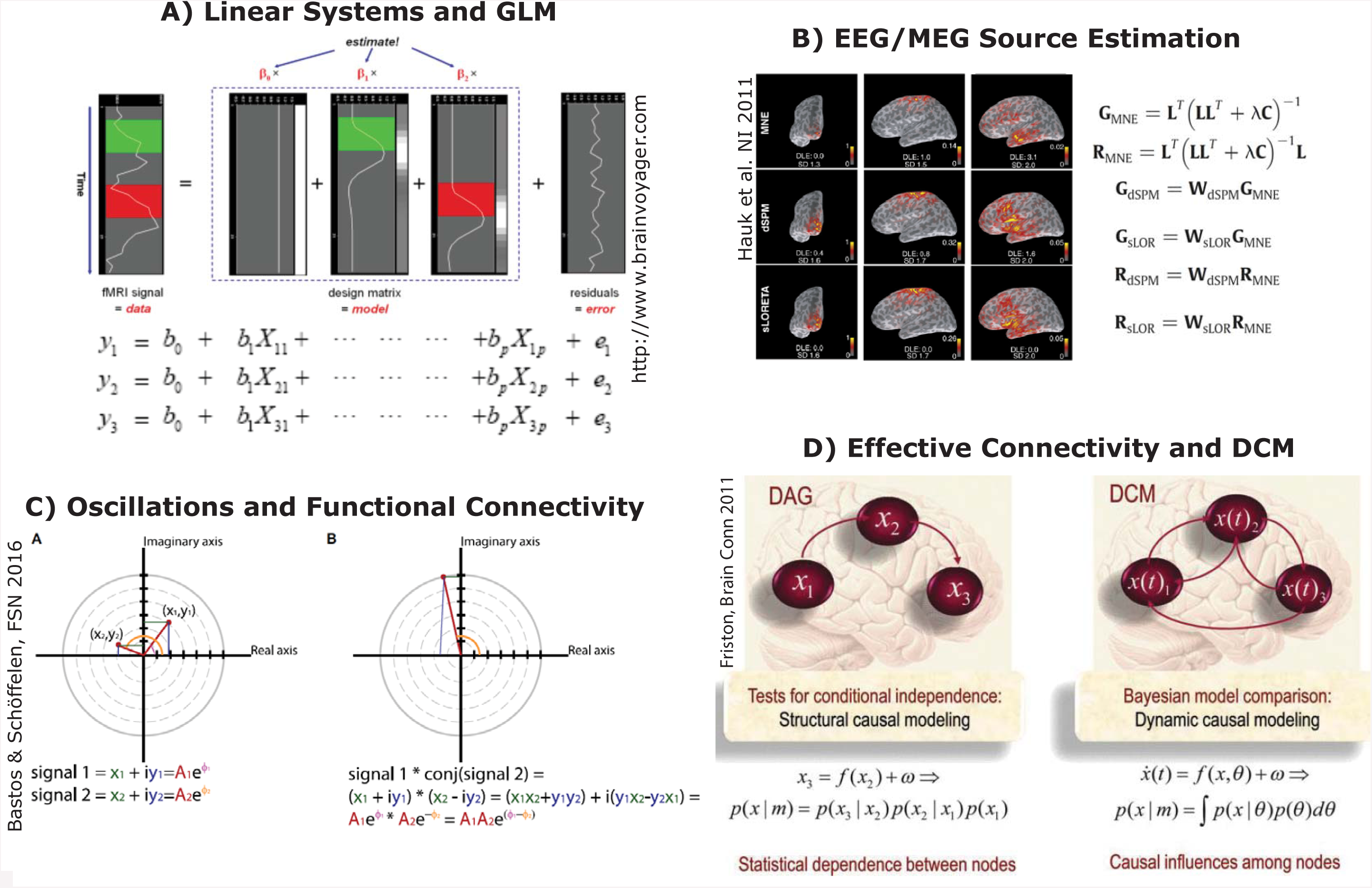
Examples of methods skills involved in the analysis and interpretation of cognitive neuroscience data. A) The general linear model, based on linear algebra (e.g. matrix multiplication and inversion), is the basis for most statistical analyses of fMRI and EEG/MEG (from http://www.brainvoyager.com). B) The EEG/MEG inverse problems, and a number of its solutions and methods to evaluate spatial resolution, are based on linear algebra (e.g. matrix multiplication and inversion) (Hauk, Wakeman, & Henson, 2011). C) Fourier and time-frequency analyses as well as spectral functional connectivity metrics are based on trigonometry and complex numbers (e.g. sines, cosines, polar representation of complex numbers) (Bastos & Schoffelen, 2015). D) Dynamic biophysical models of brain function and Bayesian model estimation, as in dynamic causal modelling (DCM), are based on calculus (e.g. integration and differentiation) (Friston, 2011).

The purpose of this survey was to estimate the current skills level in the field, as a starting point for an evidence-based discussion of current skills-levels in cognitive neuroscience. We hope that this will usefully inform the development of future skills-oriented training opportunities.

## Methods

### Participants

The survey was set up on the SurveyMonkey web-site (https://www.surveymonkey.com/, San Mateo, USA), and was advertised via neuroimaging software mailing lists and posts on the MRC Cognition and Brain Sciences Unit’s Wiki pages. It was first advertised in January 2015, and most responses (about 90%) were collected in 2015. Participation was voluntary and no monetary or other material reward was offered. Participants were informed about the purpose and nature of the survey, e.g. that it should take around 10-15 minutes of their time. They were asked not to take part more than once, and to not use the internet or books to answer the questions. The study was approved by the Cambridge Psychology Research Ethics Committee.

The number of respondents in the final analysis was 305 (139 males, 166 females). They were selected from the group of all respondents as follows.

578 participants gave consent for their data to be used for analysis (by ticking a box). Among those, we only analysed data from participants who provided responses to all methods-related questions (i.e. gave a correct, error or “no idea” response). This resulted in 322 respondents. 214 respondents skipped all methods-related questions. Although it would have been interesting to analyse the demographics of these “skippers”, many of them also skipped most of the demographic questions. For example, only 66 of them disclosed their gender (31 males, 35 females), and only 59 their undergraduate degree (37 Psychology, 14 Methods, 8 Biology).

Given the wide range of undergraduate degrees of our respondents, and the relatively small group of respondents for some of them, we grouped them into three broader categories. “Psychology” contained those who responded that their undergraduate degrees were in “psychology”, “cognitive science” and “cognitive neuroscience”. “Methods” included those who responded “physics”, “math”, “computer science”, or “biomedical engineering”. “Biology” summarised those who responded “biology” or “medicine”. For 39 respondents who indicated “Other” (e.g. “artificial intelligence”, “physiology”, “linguistics”), an appropriate group assignment was made by hand. For another 17 respondents such an assignment could not be made (e.g. musicology), and they were removed from the analysis, reducing the number of respondents in the final analysis to 305.

48% of the remaining respondents were located in the United Kingdom at the time of the survey (n=145, 83 of which from Cambridge), followed by other European countries (68, 22%), the United States of America (n=33) and Canada (5), Asia (12), Australia (4) and South America (3). 35 respondents did not provide clear information about their location.

Figure 2 shows the number of participants broken down by gender (A), current researcher status and gender (B), type of undergraduate degree and gender (C), mean age of respondents according to type of undergraduate degree and gender (D), self-reported future areas of research by gender (in percent, E), and self-rated expertise level according to undergraduate degree (in percent, F). We had more female (n=166) than male (139) respondents. Most respondents were PhD students (142), followed by post-doctoral researchers (90), undergraduate students (42) and research assistants (7). The mean age of all respondents was 29.6 (SD 6.9) years (males: 30.7 (7.5), females: 28.7 (6.2)), and varied only slightly with respect to undergraduate degree and gender. Most respondents expected to continue working in cognitive science (31%), clinical neuroscience (23%) or cognitive neuroscience (16%), with small differences between males and females. The majority rated themselves as “no experts” or “sort of” experts, though this varied with undergraduate degree. The Biology group rated themselves mostly as “no experts” (57%, compared to 22% “sort of expert” and 20% “expert”). The pattern was similar for the Psychology group, although with a smaller difference between “sort of expert” (39%) and “no expert” (46%, and 13% “expert”). The Methods group consisted mostly of “sort of experts” (48%) and “no experts” (40%), compared to 9% “experts”.

**Figure 2:**
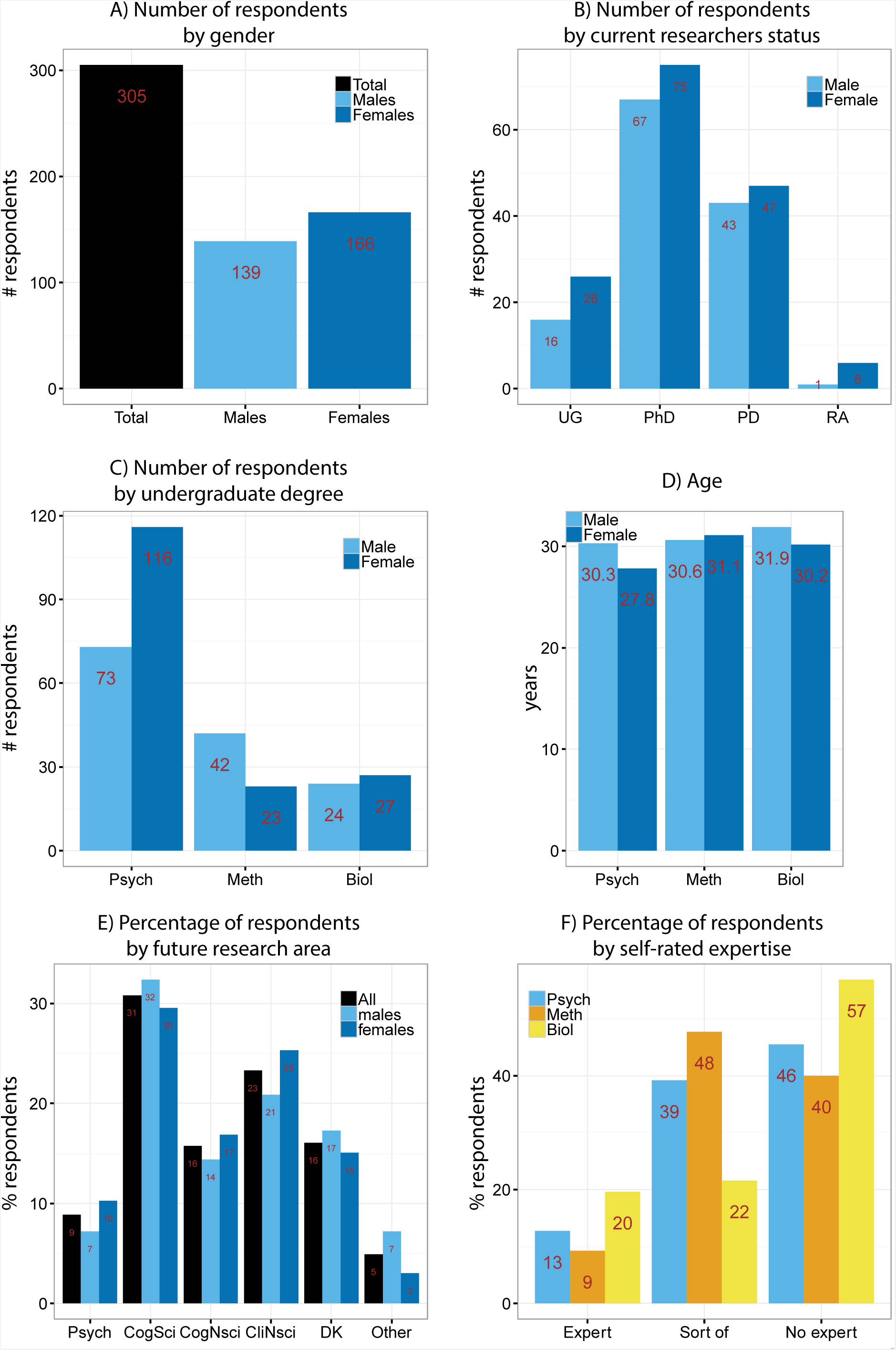
Demographics of survey participants. A) Overall number of participants and grouped by gender. B) Number of participants grouped by current researcher status and gender (UG: undergraduate student, PD: post-doctoral researcher, RA: research assistant). C) Number of participants grouped by undergraduate degree and gender (Psych: undergraduate degree in psychology-related subjects, Meth: methods-related subjects, Biol: biology-related subjects). D) Average age by undergraduate degree and gender. E) Percentage of participants depending on planned future research area and gender (Psych: future research planned in psychology, CogSci: cognitive science, CogNsci: cognitive neuroscience, CliNsci: clinical neuroscience, DK: don’t know). F) Percentage of participants who rated themselves as methods “expert”, “sort of expert” and “no expert”, according to undergraduate degree.

### The survey

The survey started with questions about demographics, such as gender, age, undergraduate degree etc. (see https://www.surveymonkey.com/r/3JL2CZX for original survey). This was followed by 18 methods-related questions (see Appendix). Questions in the survey were chosen to cover basic aspects of data acquisition and analysis in the cognitive neurosciences, especially neuroimaging (Figure 1). Questions were subjectively grouped by the author into domains of signal analysis (n=5), scientific computing (n=3), linear algebra (n=2), calculus (n=3), statistics (n=3), and physics (n=2). In brief, statistics questions probed knowledge on correlation, power analysis and the multiple comparisons problem; signal processing probed the signal-to-noise ratio, Fourier analysis, frequency spectra, complex numbers, trigonometry; calculus probed derivatives, integrals and algebraic equations; linear algebra probed vector orthogonality and vector multiplication; scientific computing probed knowledge about Linux, “for loops”, and integer numbers; physics probed Ohm’s law and electric fields. Questions were presented in multiple-choice form, with four possible answers plus a “no idea” option.

The choice of questions was constrained by

- the limited amount of time the voluntary participants were expected to invest in this survey (10-15 minutes),
- the difficulty level of the questions, which should neither bore away experts nor scare away non-experts,
- the relevance to cognitive neuroscience.

The inclusion of particular questions, based on their relevance to cognitive neuroscience research was further determined through local discussions with researchers involved in methods training, as well as by consulting text books.

### Data Analysis

Data were exported from the SurveyMonkey web-site, and converted to an Excel spreadsheet using Matlab. They were then further processed in the Software package R (Version 3.1.3, https://cran.r-project.org/). Only data from respondents who responded to every methods question were included in the analysis. For demographic questions, the total number of respondents within each category are reported. For methods skills questions, analyses focus on correct responses unless indicated otherwise. We considered the number of error and no-idea responses too low to allow meaningful interpretation of their differences. Considering the number of respondents per group (Figure 2), we think that breaking down our results by up to two factors (e.g. undergraduate degree and gender) is appropriate. The R scripts used for data analysis are available on https://github.com/olafhauk/MethodsSkillsSurvey. Data are available on request.

Our conclusions with respect to methods skills are based on the percentage of correct responses in the respective respondent groups, rather than measures of significance. A significant but small (e.g. 1%) effect would not have strong practical implications.

Nevertheless, we ran ordered logistic regression analysis with the following simultaneous factors: undergraduate degree, current researcher status, and gender. We used the function polr() from the R (i386 3.1.3) package MASS, and assessed significance using the function pnorm(). This analysis was run for overall performance as well as for sub-groups of methods questions as described below.

## Results

As described in the Methods section, our conclusions are mostly based on the mean percentages of correct responses, rather than statistical measures. Not surprisingly with the large number of participants, the results for all factors in our ordered logistic regression analysis of overall performance reached significance (p<0.05). For the six sub-groups of questions, all results for the factor Undergraduate Degree were significant except for the questions about Statistics (p=0.29), for the factor Gender except for Physics and Statistics (both p>0.8), and for the factor Researcher Status except for Linear Algebra (p=0.21) and Calculus (p=0.24). In the following, we highlight results if they consist of differences in performance of about 5% or more. The practical relevance of these effects is discussed in the discussion section.

### Overall performance

Performance across all methods questions is summarized in Figure 3A. The figure shows percentages of correct, error and “no idea” responses, respectively. Averaged across all methods questions, 72% of responses were correct, 12% error and 16% “no idea”. These results show that overall performance was not at floor or ceiling, and thus our survey should be sufficiently sensitive to be informative about differences among respondent groups.

**Figure 3:**
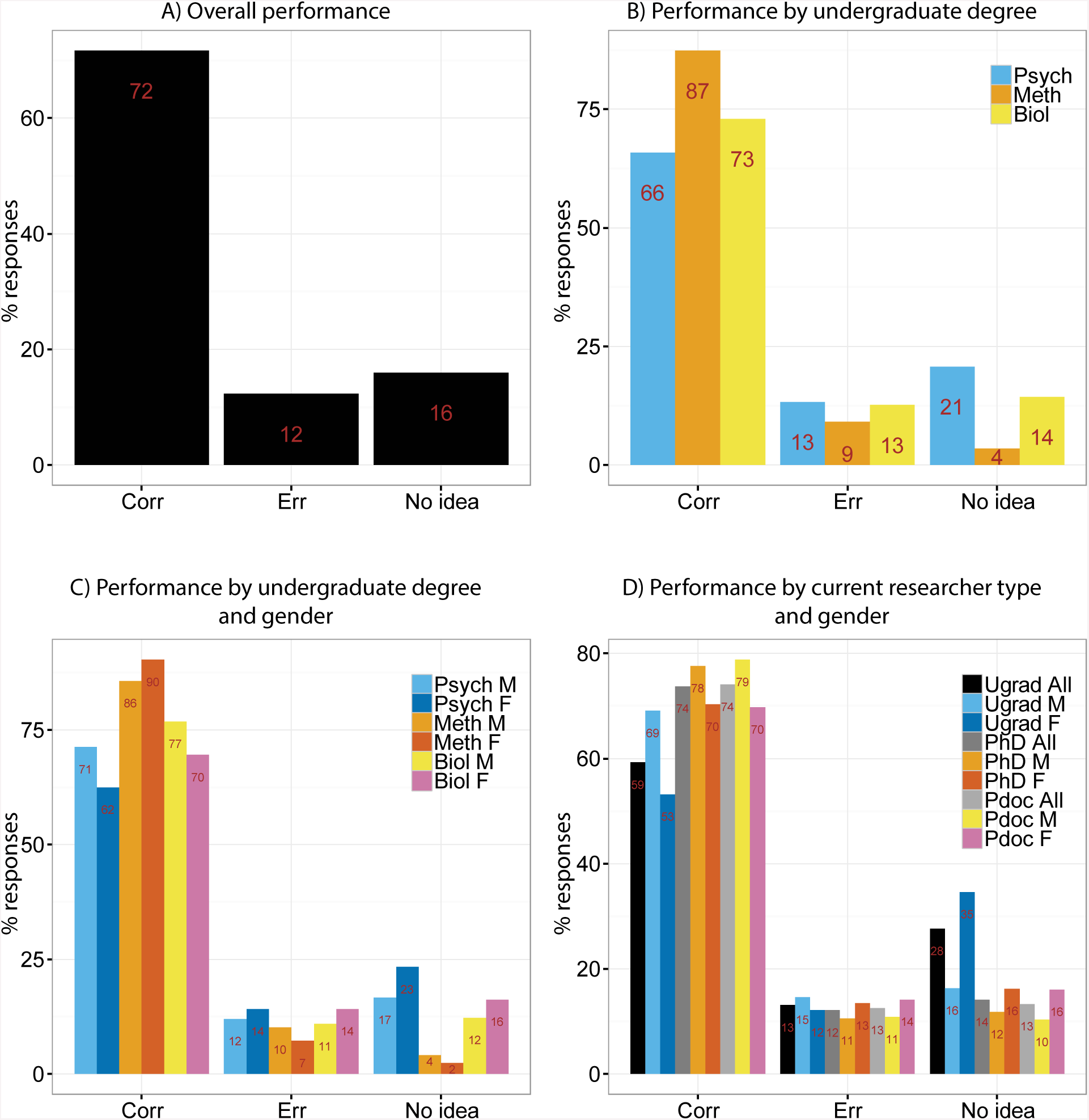
Performance across all methods questions. A) Performance across all respondents (Corr: Correct responses, Err: Incorrect responses). B) Performance broken down by undergraduate degree (Psych: Psychology subjects, Meth: Methods subjects, Biol: Biology subjects). C) Performance broken down by undergraduate degree and gender (M: Male, F: Female). D) Performance broken down by current researcher status and gender (Ugrad: undergraduate students, Pdoc: Post-doctoral researchers).

### Undergraduate degree

It is to be expected that the subject of undergraduate study strongly affects the skills being assessed in the methods questions in this survey. As described above, participants were divided into three broad undergraduate degree groupings: Psychology, Methods and Biology. The results for these groups are presented in Figure 3B. Not surprisingly, respondents with Methods undergraduate degrees provided the highest number of correct responses (87%), followed by those with Biology (73%) and Psychology (66%) undergraduate degrees.

Because the proportion of males to females differed across these groups, we also show these results broken down by gender in Figure 3C. For Psychology and Biology undergraduates, males performed slightly better than females (71% vs 62% and 77% vs 70% correct, respectively), and vice versa for Methods undergraduates (86% vs 90%).

### Current degree

In Figure 3D we present results depending on current researcher status (undergraduate/Master’s student, PhD student and post-doctoral researcher). Interestingly, there is some improvement from the undergraduate (59%) to the PhD (74%) and post-doctoral (74%) level, but no improvement from PhD to post-doctoral level. Males showed better performance in all three groups (undergraduates: 69% vs 53% PhD: 78% vs 70%, Post-doc: 79% vs 70%). This result is likely to be confounded by undergraduate degree (Figure 2C), but it shows that gender differences in methods skills persist at higher academic degrees.

### Self-rated expertise

Though not all cognitive neuroscientists need methods skills at the same level, it is nonetheless important that students and post-doctoral researchers have a realistic view of their own methods skills. We therefore asked participants to rate themselves as “Expert”, “Sort of expert” or “No expert” (see also Figure 2F). We present our results broken down into these categories in Figure 4A. Performance differs surprisingly little between experts and sort-of experts (81% vs 75%), but there is a bigger gap to the non-experts (66%). The results are further broken down by undergraduate degree in Figure 4B. In the Methods group, performance was generally high with small differences among expertise levels (“expert”: 92%, “sort of expert”: 88%, “no expert”: 86%). This gradient was somewhat steeper in the Biology group (83%, 79%, 71%), and largest for Psychology (79%, 70%, 58%). Respondents from the methods group who rated themselves as “no experts” still performed better than experts from the other two groups.

**Figure 4:**
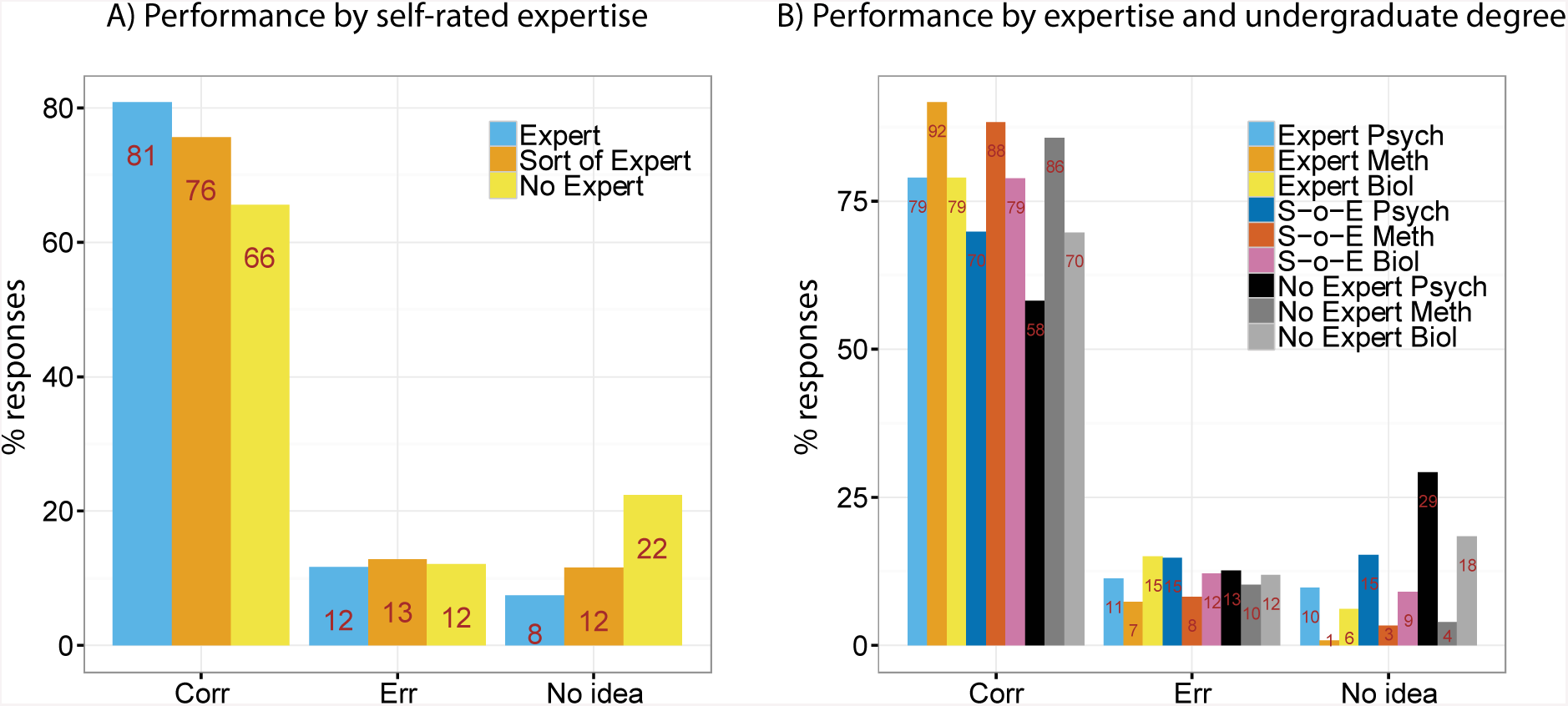
Performance by self-rated methods expertise level (A) and expertise by undergraduate degree (B).

### Sub-groups of Methods Questions

Our questions covered different aspects of neuroimaging data acquisition and analysis. We grouped our questions into six categories that reflect important aspects of neuroimaging data analysis, i.e. Linear Algebra, Signal Analysis, Calculus, Programming, Physics and Statistics. Results for these groups are presented in Figure 5. Because in our previous analyses the largest differences occurred with respect to undergraduate degree, results were split according to these groups.

**Figure 5:**
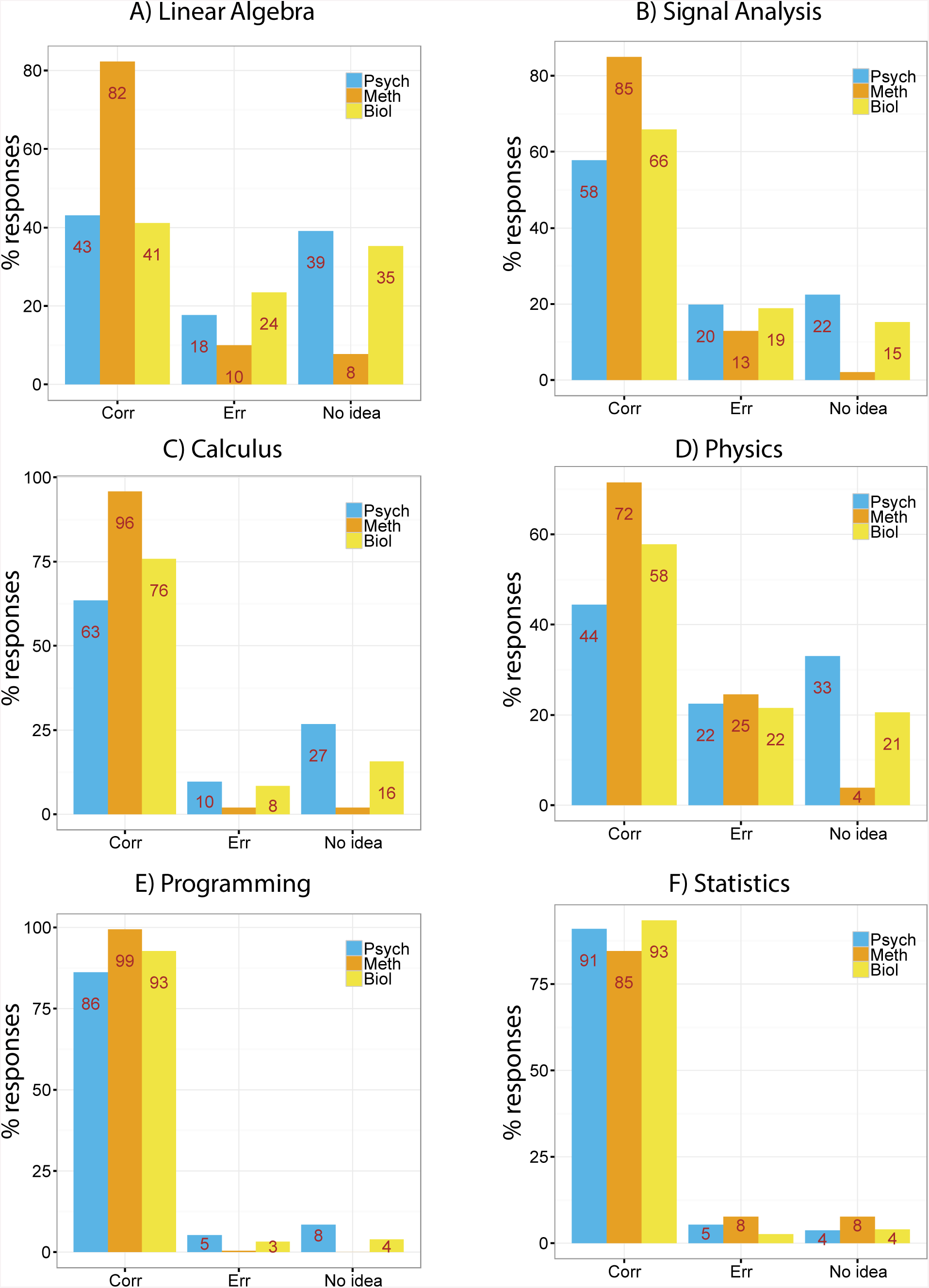
Performance for specific question groups by undergraduate degree. A) Linear Algebra (questions about vector orthogonality and vector multiplication). B) Signal Analysis (signal-to-noise ratio, Fourier analysis, frequency spectra, complex numbers, trigonometry). C) Calculus (derivatives, integrals and algebraic equations). D) Programming (Linux, “for loops”, integer numbers). E) Physics (Ohm’s law, electric field and potential). F) Statistics (correlation, power analysis, multiple comparisons problem).

Differences among groups were most striking for Linear Algebra (Figure 5A), where performance for Psychology and Biology undergraduate degree holders was only 43% and 41% correct respectively, compared to 82% for Methods undergraduates. Similar results were obtained for Signal Analysis (5B, 58% and 66% compared to 85%), and Calculus (5C, 63% and 76% compared to 96%). A similar pattern, although at generally lower performance, was observed for Physics (5D, 44% and 58% compared to 72%). For Programming (5E) performance was more equally distributed (86% and 93% compared to 99%), and for Statistics (5F) the pattern was reversed (91% and 93% compared to 85%).

### Demand for more skills-oriented training

Finally, we asked participants whether they would like to receive more training on the methods topics covered by this survey. The results are presented in Figure 6. Figure 6A shows that the majority (63%) of participants would like to receive “a lot” or “significantly” more training on these topics, with an additional 30% asking for “a little” more training. Only 4% responded that they do not want more training at all, and another 3% did not know. There was more demand among female (31%) compared to male (24%) respondents for “a lot” more training. Figures 6B-C break these results down further, and show that the general pattern is the same for different researcher types (Fig. 6B) and undergraduate groups (Fig. 6C). Methods undergraduate degree holders have lower demand for methods training than their Psychology and Biology counterparts, but most of them still vote for “a little” more training, and about 50% still for “a lot” or “significantly” more.

**Figure 6:**
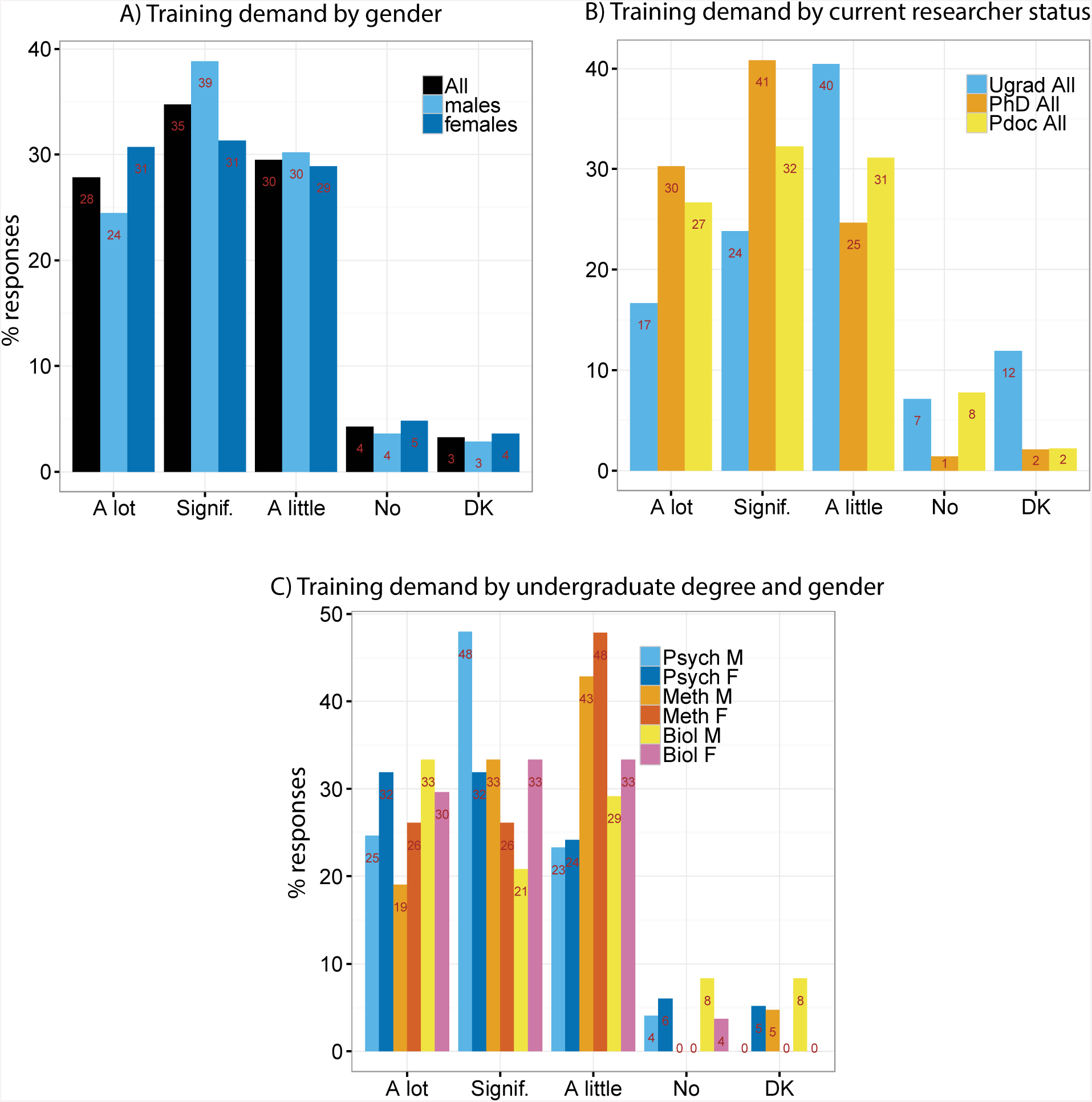
Demand for more training on topics covered in this survey for all participants and by gender (A), current researcher status, (B) as well as undergraduate degree and gender (C). Signif.: Significantly. DK: Don’t know.

## Discussion

We performed an online survey among 305 mostly junior scientists in the cognitive neurosciences, and evaluated their basic methods skills in the areas of signal processing, linear algebra, calculus, statistics, programming, and physics. The topics of this survey covered basic textbook knowledge relevant to the analysis of neuroimaging data and computational modelling, as well as to the interpretation of results in the cognitive neuroscience literature. Our results suggest that there is room for improvement with respect to the methods skills among junior researchers in cognitive neuroscience, that researchers are aware of this, and that there is strong demand for more skills-oriented training opportunities.

Performance varied with respect to undergraduate degree, current researcher status and gender. Overall, performance was at 72% correct, i.e. neither at bottom nor at ceiling. It is difficult to interpret this overall performance level without knowing more about the motivation and response style of participants. For example, it is impossible to determine for certain whether those who did not complete this survey did so because it was too challenging or too boring or irrelevant to them, which could potentially have biased our results. This is a problem with many surveys, not just online. However, performance was high where expected, e.g. for basic questions about programming (“What is Linux?”, “What is a for-loop?”), and the way the pattern of results related to undergraduate degree was very plausible (respondents with a methods undergraduate degrees performing better than those with psychology or biology degrees). Most participants were recruited through e-mail announcements on software mailing lists, and it is likely that they noticed these e-mails because they are actively engaged in neuroimaging projects. Most participants indicated that they would like to receive significantly or a lot more training on the topics covered by this survey. This suggests that the participants of this survey took an interest in the subject and were motivated to perform well. Thus, we conclude that the group differences discussed below are informative about junior researchers in the cognitive neurosciences.

Not surprisingly, we found the most striking differences in performance among respondents with a Methods undergraduate degree versus those with a Psychology or Biology degree (note that we grouped different undergraduate degrees together). Overall performance of the Psychology group was about 20% lower than the Methods group, with Biology in-between. This difference was of similar magnitude for questions on signal processing, and was much larger (almost 50%) for questions on linear algebra. These two topics are fundamental to the analysis and interpretation of neuroimaging data. The particular questions addressed common problems in neuroimaging analysis, such as the orthogonality of vectors and the interpretation of a frequency spectrum. The multiplication of vectors and matrices is central to the general linear model (GLM), which is ubiquitous in fMRI and EEG/MEG data analysis (e.g. https://en.wikibooks.org/wiki/SPM). On a practical level, understanding why vectors and matrices of certain dimensions cannot be multiplied with each other can help with the debugging of analysis scripts (e.g. “matrix dimensions do not agree” in Matlab, Python or R). For example, a lack of awareness that for matrices the commutative law does not necessarily hold (i.e. that **A**\***B** is not necessarily the same as **B**\***A**, unlike for scalar numbers), can potentially lead to incorrect results. In the domain of signal analysis, familiarity with the basic concepts of frequency spectra and Fourier analysis is a pre-requisite for time-frequency (e.g. using wavelets) and spectral connectivity (e.g. using coherence) analyses (e.g. Bastos & Schoffelen, 2015). Calculus (e.g. differentiation and integration) is required to deal with differential equations, which are the basis for dynamic causal modelling and Bayesian inference (Friston, 2010; Friston, 2011).

Our findings not only have implications for data analysis, but also theory development. Interest in computational modelling of cognition in combination with neuroimaging has grown over the last decade or so (Friston & Dolan, 2010; Kriegeskorte, 2015; Turner, van Maanen, & Forstmann, 2015). It is often said that the brain is the most complex information processing system we know. It would be puzzling if studying this complex information processing system required less methods skill than investigating “simple” radios or microprocessors (Jonas & Kording, 2017; Lazebnik, 2002). If cognitive neuroscience is to become a quantitative natural science, then cognitive neuroscientists need quantitative skills.

Our results should also be taken into account by methods developers and tutors. Methods should be described using concepts and terminology that the target audience understand. If a paper starts with the most general theoretical description of a method in order to then derive special cases that are relevant for practical purposes, many respondents of this survey will be lost straightaway. For example, starting with Bayesian model estimation in order to derive ordinary linear least-squares regression may not produce the desired learning outcome for researchers from a psychology or biology background. It may be better to try it the other way round: start with a problem the target audience are familiar with, and then address its limitations and how to look at the problem at increasingly general levels.

Interestingly, we found an increase in performance from the undergraduate to the PhD and post-doctoral level, but no difference between the latter two. This suggests that researchers maintain the skills level they have acquired during their undergraduate and PhD degrees. This is fine if they continue the kind of work they were trained for during the undergraduate or PhD projects. However, this is unlikely to be the case for many researchers. Firstly, cognitive neuroscience is a fast developing field where methodological innovations are a strong driving force. It is highly likely that researchers require new methods skills during their PhD or post-doctoral work. Secondly, it is hard to conceive that students already know precisely what kind of work they will do at the post-doctoral stage. They may start with an interest in behavioural research or cognitive theory, but later discover their passion for brain imaging. A lack of skills-oriented training opportunities makes a change in research area much harder and frustrating, if not impossible. This is particularly true for an interdisciplinary and dynamic research area such as cognitive neuroscience.

In order to choose the right type of training, or the right research topic to work on, it is important to know your own strengths and weaknesses if it comes to methods skills. We asked our respondents to rate themselves as “experts”, “sort of experts”, and “no experts”. Interestingly, only 8% in the Methods undergraduate group rated themselves as experts, less than for Psychology and Biology (who led the table with 19%). However, “no expert” Methods respondents still outperformed all other non-methods groups. This suggests that researchers estimate their own skill level relative to members of their own group. Biologists and psychologists who work in neuroimaging may have more experience with data analysis than their peers who work in other domains, but still less than researchers from an engineering or physics background. The latter, in turn, may think that because they are not working in engineering or physics anymore, their skill level is lower, even though it may still be higher than for many other researchers in their field. This discrepancy may lead to a lack of awareness of one’s own limitation and the need for advice on the one hand, and an undervaluation of one’s own capacity to provide training and advice to peers on the other. It is important to know what you know and what you don’t know. Not everyone needs or wants to be a methods expert. But those who do will need to spend a significant amount of time and effort on appropriate training. Those who don’t should know where to get advice when needed.

We also found overall performance differences with respect to gender, albeit these effects were smaller than those among undergraduate groups. Performance was higher for males than females for all respondent groups except for Methods undergraduates where the effect was reversed. We can only speculate about the reasons. Women are less likely to choose methods-related subjects at school or at the undergraduate level (Stoet & Geary, 2018). During their PhD or post-doctoral research, women may feel less encouraged to develop methods skills because methods development and scientific computing are currently dominated by males. Our finding that women outperformed men in the Methods undergraduate group demonstrates the obvious, namely that both genders can achieve similar skill levels when given the same opportunities. It has been demonstrated that while girls outperform boys in science subjects (and others) at school, they are underrepresented in STEM degrees, which may be related to their attitude towards science and their self-estimation of academic skills (Stoet & Geary, 2018). The superior performance of women with Methods undergraduate degrees in our survey may reflect the fact that they are in the minority in this community, and are more motivated or feel greater pressure to succeed. Skills-oriented training programmes may therefore contribute to equal opportunities in cognitive neuroscience. Importantly, women should engage in tutoring and teaching methods-related skills, as increasing the visibility of women engaging in methods research can affect perception and feelings of belonging or inclusion of more junior members of the research community.

We found the largest differences between groups of respondents with different undergraduate degrees. This suggests that researchers from psychological and biological backgrounds are not necessarily entering the field of cognitive neuroscience with the skills required to analyse data and meaningfully interpret cognitive neuroscience findings reported in the literature. It is possible that small differences at early career stages, if not corrected, may amplify over time. Researchers who get frustrated with methodological challenges may decide to change fields. This may also be relevant for the gender differences discussed above. It is therefore important to provide opportunities for researchers at different stages, especially at PhD and junior post-doctoral level, to develop and maintain their methods skills.

Biologists and medical students undergo basic training in maths and physics during their undergraduate degrees, but at that time may not have specialised in any particular subject and may not be aware of the importance of these subjects for their future work. Psychologists often receive training in statistics, but not in signal processing, linear algebra or physics. These issues are addressed to some degree in cognitive neuroscience programmes that are becoming increasingly popular. However, for researchers who move into neuroimaging after their undergraduate or post-graduate degrees, this may be too late. We therefore recommend that the relevance of these topics is clearly communicated to students and post-doctoral researchers as well as supervisors.

The necessary skills for a research project should be identified both by supervisors and students or post-doctoral researchers. Our survey was dominated by respondents from the United Kingdom (48%) and Europe (22%). Skills levels may vary depending on country and institution, and ideally should be evaluated locally before students enter a project. Gaps in the skills set can then be addressed by local training programmes, which can be integrated with courses on cognitive or clinical neuroscience, as well as by external workshops or online resources. While a range of software-related workshops exist that focus on the “doing” part of data analysis, our survey results indicate that more needs to be done about the “understanding” part. In the future, the cognitive neuroscience community could create guidelines about a “core skills set” for cognitive neuroscientists, taking into account different sub-disciplines such as neuroimaging, computational modelling and clinical neuroscience.

Good teaching, whether on methods or other subjects, should be valued and appreciated (as lamented previously, e.g. Anonymous Academic (2014)). This needs to be budgeted for both in terms of money and time. As a bonus, methods skills are highly transferrable and can be useful in careers outside academia.

We hope that our survey, despite its limitations, provides a starting point for further evidence-based discussion about the way cognitive neuroscientists are trained.

## Acknowledgments

We would like to thank Fionnuala Murphy, Johan Carlin and Alessio Basti for her very helpful comments and corrections on a previous version of this manuscript, Rogier Kievit for glancing over my R code (but he should remain unblamed for any incorrectly placed brackets or hyphens), and members of the Cognition and Brain Sciences Unit who have engaged in numerous discussions about training-related issues over the last few years. We are grateful for the funding we received from the Medical Research Council UK (SUAG/019 RG91365).

## Appendix: Survey questions and multiple-choice answers

The methods-related questions of the survey, grouped into categories, are listed below. Every question also had a “No idea” option, which is not listed here.

## Statistics

The correlation coefficient between two signals describes

- whether one signal is larger than the other
- the linear relationship between the signals
- whether one signal is the derivative of the other
- the sound quality of the signals

In statistics, a power analysis can be used to determine

- whether an analysis is sensitive enough to detect an effect of a certain size
- how much computer memory you need to perform an analysis
- whether an effect is statistically significant
- how strong your signal is

The statistical “multiple comparisons problem” refers to

- the decrease of statistical power with large data sets
- the increased uncertainty in your result with too few samples
- the increase of computing power required to perform multiple statistical tests
- the increased likelihood of false positives if one performs multiple independent tests

## Signal Analysis

The “signal to noise ratio” (SNR) refers to

- a measure for the amplitude of the desired part of the signal divided by a similar measure for the undesired part of the signal
- a measure for the amplitude of the undesired part of the signal divided by a similar measure for the desired part of the signal
- the ratio between two random signals
- a statistical comparison between a structured and a random model

If you have two signals with frequencies f1 and f2, then the sum of these signals has peaks in the frequency spectrum at which frequency or frequencies?

- f1+f2 and f1-f2
- f1 and f2
- f1*f2
- (f1+f2)/2

A Fourier analysis is

- a method to decompose signals into shorter sequences
- a method to determine the four most significant components of a signal
- a method to decompose signals into complex polynomials
- a method to decompose signals into sine and cosine functions

The sine of an angle in a right-angled triangle is equal to

- opposite divided by adjacent leg
- adjacent divided by opposite leg
- opposite leg divided by hypotenuse
- adjacent leg divided by hypotenuse

A complex number is

- a number that cannot be expressed as the ratio of two integers
- a number that is difficult to keep in working memory
- a number that involves the square root of −1
- a number with more than 10 digits

## Calculus

If function f is the integral of the function g, then

- f divided by g is unity
- the derivative of f is g
- the integral of f is g
- f and g are orthogonal

The derivative of “2 times x-squared” is:

- 4 times x
- 2 times x-squared
- 4 times x-cubed
- 2 times x

What is the solution to 3*x – 2 = 7?

- 1
- 3
- 5
- 12

## Linear Algebra

What is the (scalar) product of the two vectors [1 2] and [3 4]?

- 4
- 21
- 10
- 11

Which of these pairs of vectors are orthogonal?

- [1 2], [1 2]
- [0 1], [0 1]
- [1 2], [3 4]
- [1 −2], [2 1]

## Programming

An integer number is

- a number that can be written without a fractional component
- a number that cannot be divided
- a prime number smaller than 100
- a positive number

In software programming, a “for loop” is a statement that

- circumvents an error message
- asks for further information
- allows a piece of code to be repeatedly executed
- lets the program return to a previous statement

“Linux” is

- the first scientific computer game
- a powerful text editor
- a software for scientific data analysis
- an open source computer operating system

## Physics

Ohm’s law, i.e. the relationship between voltage and current in an electric circuit, states that

- Voltage = Resistance / Current
- Current = Resistance * Voltage
- Current = Voltage / Resistance
- Current = Voltage - Resistance

The electric field is

- the gradient of the electric potential
- the integral of the electric potential
- the square of the electric potential
- the distribution of the electric potential visualised as arrows

